# Mitochondrial UPR negatively regulates wounding-induced innate immune response in *C. elegans* epidermis

**DOI:** 10.1101/2021.09.19.460990

**Authors:** Jianzhi Zhao, Hongying Fu, Hengda Zhou, Xuecong Ren, Yuanyuan Wang, Suhong Xu

## Abstract

Tissue damage elicits a rapid innate immune response that is essential for efficient wound healing and survival of metazoans. It is well known that p38 MAPK kinase, TGF-β, and hemidesmosome signaling pathways have been involved in wounding-induced innate immunity in *C. elegans*. Here, we find that loss of function of ATFS-1 increased innate immune response while an elevated level of mitochondrial unfolded protein response (mitoUPR) inhibits the innate immune response upon epidermal wounding. Epidermal wounding triggers the nucleus export of ATFS-1 and inhibits themitoUPR in *C. elegans* epidermis. Moreover, genetic analysis suggests that ATFS-1 functions upstream of the p38 MAP kinase, TGF-β and DAF-16 signaling pathways in regulating AMPs induction. Thus, our results suggest that the mitoUPR function as an intracellular signal required to fine-tune innate immune response after tissue damage.

**Highlights:**

— ATFS-1 inhibits the wounding-induced innate immune response in *C. elegans*
— mitoUPR inhibits wounding-induced innate immune response
— Epidermal wounding induces downregulation of ATFS-1 in the epidermis
— mitoUPR acts upstream of p38MAPK and TGF-β pathways to regulate innate immunity

## INTRODUCTION

Wounding-induced immunity is a key feature for wound healing in all eukaryotic organisms. It is known that perturbations to protein synthesis, proteolysis, and mitochondrial activity are sufficient to activate innate immune responses after pathogen infection (Dunbar et al., 2012; Melo and Ruvkun, 2012; Richardson et al., 2010). *Caenorhabditis elegans* epidermis constantly encounters myriad environmental insults, including skin-puncturing pathogens and mechanical damage, which can break the epidermal barrier and activate self-defense responses by elevating the production of antimicrobial peptides (AMPs) (Pujol et al., 2008; Xu and Chisholm, 2011). The damage to the skin triggers a rapid innate immune response by inducing multiple AMPs such as neuropeptide-like proteins (NLPs) and caenacins (CNCs) to defend against sterile infection (Pujol et al., 2008; Zugasti and Ewbank, 2009), and this process is regulated by p38 MAP kinase, TGF-β, and hemidesmosome signaling pathways (Kim et al., 2002; Pujol et al., 2008; Zhang et al., 2015; Zugasti and Ewbank, 2009).

Cells respond to mitochondrial dysfunction by activating the mitochondrial unfolded protein response (mitoUPR) to detect pathogenic or toxic bacteria by monitoring mitochondrial function and initiating an innate immune response accordingly (Liu et al., 2014; Nargund et al., 2015; Nargund et al., 2012; Pellegrino et al., 2014). ATFS-1 is a key transcription factor required for the activation of mitoUPR. In healthy animals, ATFS-1 is efficiently imported into mitochondria and degraded. Under mitochondrial stress, mitochondrial import efficiency is reduced, allowing ATFS-1 to accumulate in the nucleus, where it can activate a protective transcriptional response (Haynes and Ron, 2010; Lin and Haynes, 2016; Nargund et al., 2012). A recent study reported that defective ATFS-1 makes *C. elegans* susceptible to opportunistic *P. aeruginosa*, while constitutive activation of ATFS-1 improves the scavenging capacity of this pathogen in the intestine (Pellegrino et al., 2014). However, it is still unclear whether mitoUPR involves in the tissue damage-induced innate immune response.

## RESULTS

### ATFS-1 inhibits wounding-induced innate immune response

We investigated the wounding-induced innate immunity by examining the expression of the AMPs using a transcriptional reporter strain *Pnlp-29::GFP*, which can be strongly induced upon wounding (Pujol et al., 2008). We observed that an obvious upregulation of *Pnlp-29-GFP* expression in wild-type (WT) animals 4 and 7 hours post-wounding (h.p.w.) (Figure 1A). We then examined the expression of *Pnlp-29-GFP* in *atfs-1(tm4525)* mutant, in which a larger fragment of genomic DNA was deleted and reduced the mitoUPR (Nargund et al., 2012; Pellegrino et al., 2014). The expression of *Pnlp-29-GFP* in *atfs-1(tm4525)* without wounding is similar to WT but significantly enhanced after wounding (Figure 1A), suggesting the possibility that ATFS-1 functions in the wounding-induced innate immune response.

**Figure 1.**
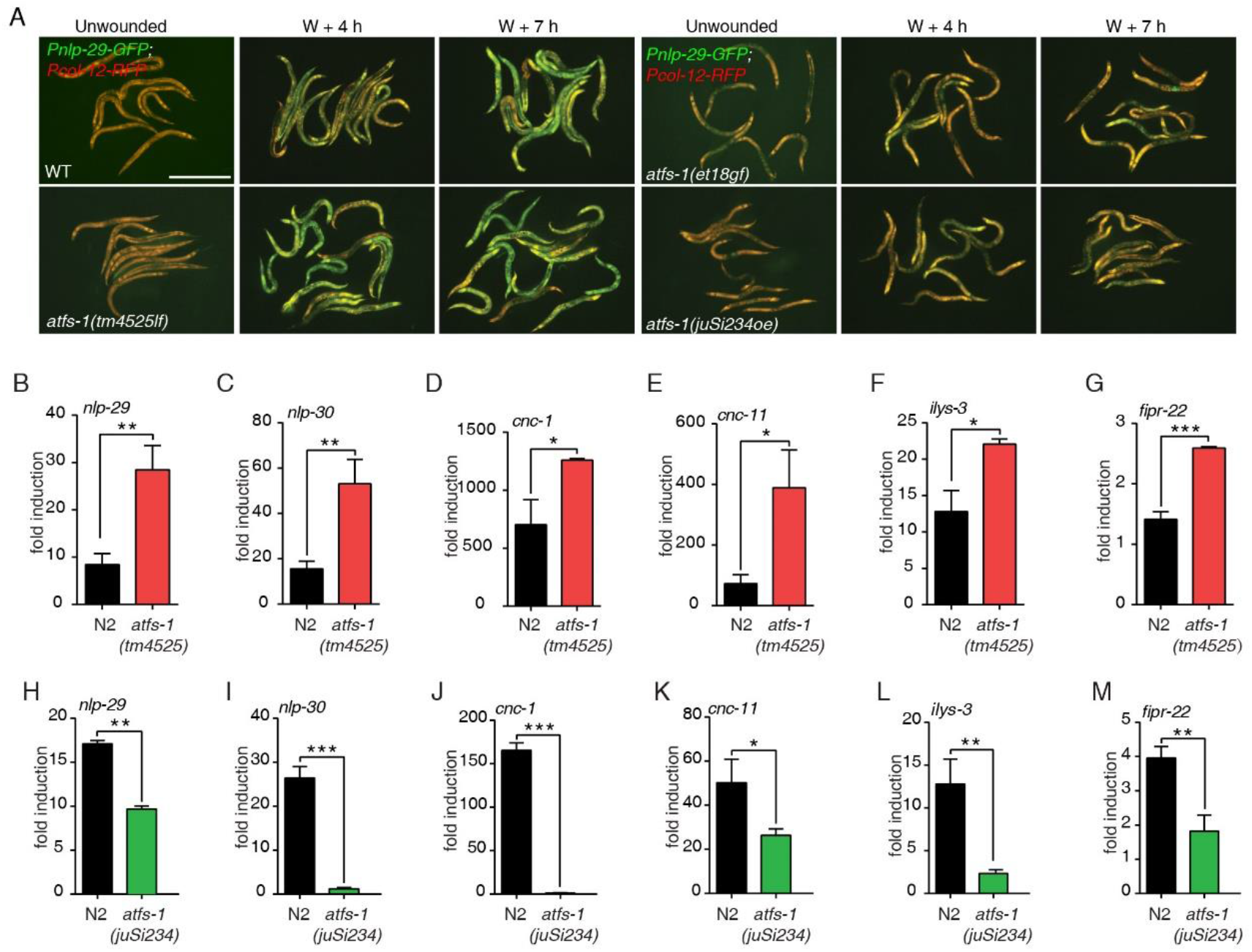
ATFS-1 inhibits epidermal wounding-induced innate immune responses. (A) The green fluorescence in transgenic worms carrying a *Pnlp-29::GFP* reporter was observed post wounding in worms generated by crossing with WT, *atfs-1* loss-of-function mutant *atfs-1(tm4525*), *atfs-1(et18gf*) or epidermis specific overexpression mutant *atfs-1(juSi234)*. Scale bars, 500 μm. (B-G) *nlp-29, nlp-30, cnc-1, cnc-4, ilys-3*, and *fipr-22* expression levels as determined by qPCR in WT and *atfs-1(tm4525)* animals. N = 3 replicates. Bars indicate the mean ± SEM. *, P < 0.05, **, P < 0.01, ***, P < 0.001 (Student’s t-test).(H-M) *nlp-29, nlp-30, cnc-1, cnc-11, ilys-3*, and *fipr-22* expression levels as determined by qPCR in WT and *atfs-1(juSi234)* animals. N = 3 replicates. Bars indicate the mean ± SEM. *, P < 0.05, **, P < 0.01, *** P < 0.001 (Student’s t-test).

To confirm this observation, we used quantitative PCR (qPCR) to measure the relative expression of other AMPs, including *nlp-29, nlp-30* (Pujol et al., 2008) and *cnc-1,cnc-11* (Zugasti and Ewbank, 2009). Upon wounding, these two types of AMPs were expressed at significantly higher levels in both *atfs-1(tm4525)* mutants compared to WT (Figure S1A). Knockdown of *atfs-1* by RNAi also enhanced the AMPs expression after wounding compared to L4440 (empty vector) control (Figure S1B). As we compared the expression change of AMPs before and after wounding, we thus quantified the AMPs expression by measuring the fold induction (Zugasti and Ewbank, 2009). Consistently, both NLPs and CNCs were significantly upregulated in *atfs-1(tm4525)* and *atfs-1(tm4919)* two loss-of-function mutants as well as RNAi knockdown animals after wounding (Figure 1B-E, Figure S1C-F). In addition, the expression levels of *fipr-22* and *ilys-3*, which are known to function in defense against fungal pathogen infections in the epidermis (Gravato-Nobre et al., 2016; Kim and Ewbank, 2018), were also significantly increased in *atfs-1* mutants and RNAi knockdown animals after wounding (Figure 1F-1G, Figure S1G-J). Collectively, these results suggest that ATFS-1 is required for negative regulation of innate immune response upon skin damage.

We then examined if ATFS-1 is sufficient to inhibit the wounding-induced innate immune responses. Gain-of-function mutation of *atfs-1(et18gf)*, a point mutation by which mitoUPR is constitutively activated (Rauthan et al., 2013), displayed a reduction in the extent of induction of *Pnlp-29-GFP* reporter (Figure 1A). Quantitative analysis *nlp-29, nlp-30, cnc-11, CNCs, ilys-3*, and *fipr-22* showed a lower level of induction after wounding (Figure S2A-E). To further confirm this observation, we expressed ATFS-1 specifically in the epidermis under the control of the *col-19* promoter. Overexpression of ATFS-1 in the epidermis significantly activated the mitoUPR (Figure S2F), and inhibited the induction of NLPs, CNCs, *ilys-3*, and *fipr-22* innate immune response genes after wounding (Figure 1H-M, Figure S2G). Together, these observations support the notion that over-expression of ATFS-1 inhibits the wounding-induced innate immune responses.

### mitoUPR inhibits wounding induced innate immunity

ATFS-1 is a key transcriptional factor required for mitoUPR (Nargund et al., 2012), we thus determine whether mitoUPR inhibits the wounding-induced innate immune responses. To this end, we knocked down *spg-7*, which encodes a mitochondrial protease required to inhibit mitoUPR (Haynes and Ron, 2010) (Figure S3A). We observed a significant inhibition of *Pnlp-29::GFP* induction 4 and 7 h.p.w. (Figure 2A). Moreover, qPCR analysis showed that knockdown of *spg-7* also inhibited the induction of *nlp-29, nlp-30, cnc-1, cnc-11, fipr-22*, and *fipr-23* after wounding (Figure 2B-I). In contrast, knockdown of *ubl-5*, which is required for mitoUPR activation (Benedetti et al., 2006; Haynes et al., 2007a; Haynes and Ron, 2010) (Figure S3A), significantly increased the induction of AMPs by wounding (Figure 2A-I). Furthermore, knockdown of *clpp-1*(RNAi), which encodes an ATP-dependent mitochondrial protease required for the induction of mitochondrial chaperone genes in response stress (Haynes et al., 2007a), increased the induction of AMPs (Figure S3B-E). Together, these results indicate that mitoUPR signal negatively regulates innate immune response after epidermal wounding.

**Figure 2.**
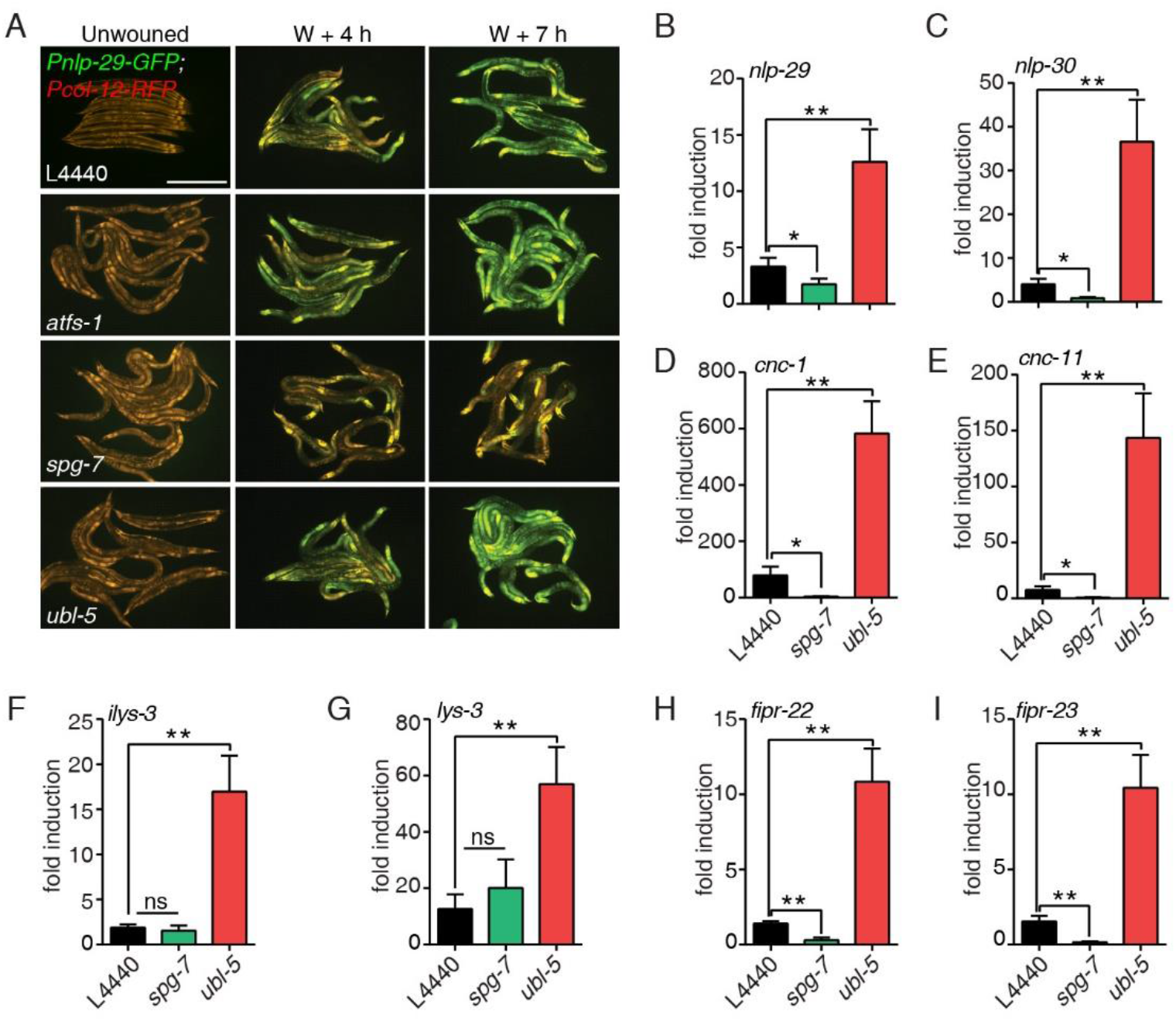
mitoUPR inhibits the induced expression innate immune responses genes upon epidermal wounding. (A) The green fluorescence in transgenic worms carrying a *Pnlp-29::GFP* reporter was observed in the wounded animals fed with an L4440 empty vector, *atfs-1, spg-7*, or *ubl-5* RNAi. Scale bars, 500 μm. (B-I) *nlp-29, nlp-30, cnc-1, cnc-11, ilys-3, lys-3, fipr-22*, and *fipr-23* expression levels as determined by qPCR in WT animals fed with an L4440 empty vector, *spg-7*, or *ubl-5* dsRNA. N = 3 replicates. Bars indicate the mean ± SEM. ns, P > 0.05, *, P < 0.05, **, P < 0.01 (Student’s t-test).

### Epidermal wounding inhibits mitochondrial unfolded protein responses

We then determine how epidermal wounding affects the activity of mitoUPR. To this end, we examined the expression of a *Phsp-6::GFP*, an indicator of mitoUPR activation in *C. elegans* (Haynes et al., 2007b). However, we did not observe an apparent increase or reduction of *Phsp-6::GFP* expression after wounding (Figure 3A, Figure S4A). As needle puncture damaged the small region of the epidermis, the live imaging system may not be able to detect the local response of mitoUPR at the protein level. To further test this, we measured the transcriptional expression of *hsp-6* and *hsp-60*, mitochondrial chaperone protein-coding genes that is activated by mitoUPR (Yoneda et al., 2004). Surprisingly, we observed a significant reduction in either *hsp-6* or *hsp-60* as early as 15 minutes to 3 hours after wounding (Figure 3B and Figure S4B), suggesting that the epidermal wounding may inhibit the mitoUPR. To further test this, we examined the expression of *Phsp-6::GFP* in animals with an elevated level of mitoUPR by ATFS-1 epidermal overexpression (Figure 3A). We observed an apparent reduction of P*hsp-6*::GFP fluorescence signal after wounding (Figure 3A, Figure S4A). Together, these results suggest that epidermal wounding inhibits mitoUPR response.

**Figure 3.**
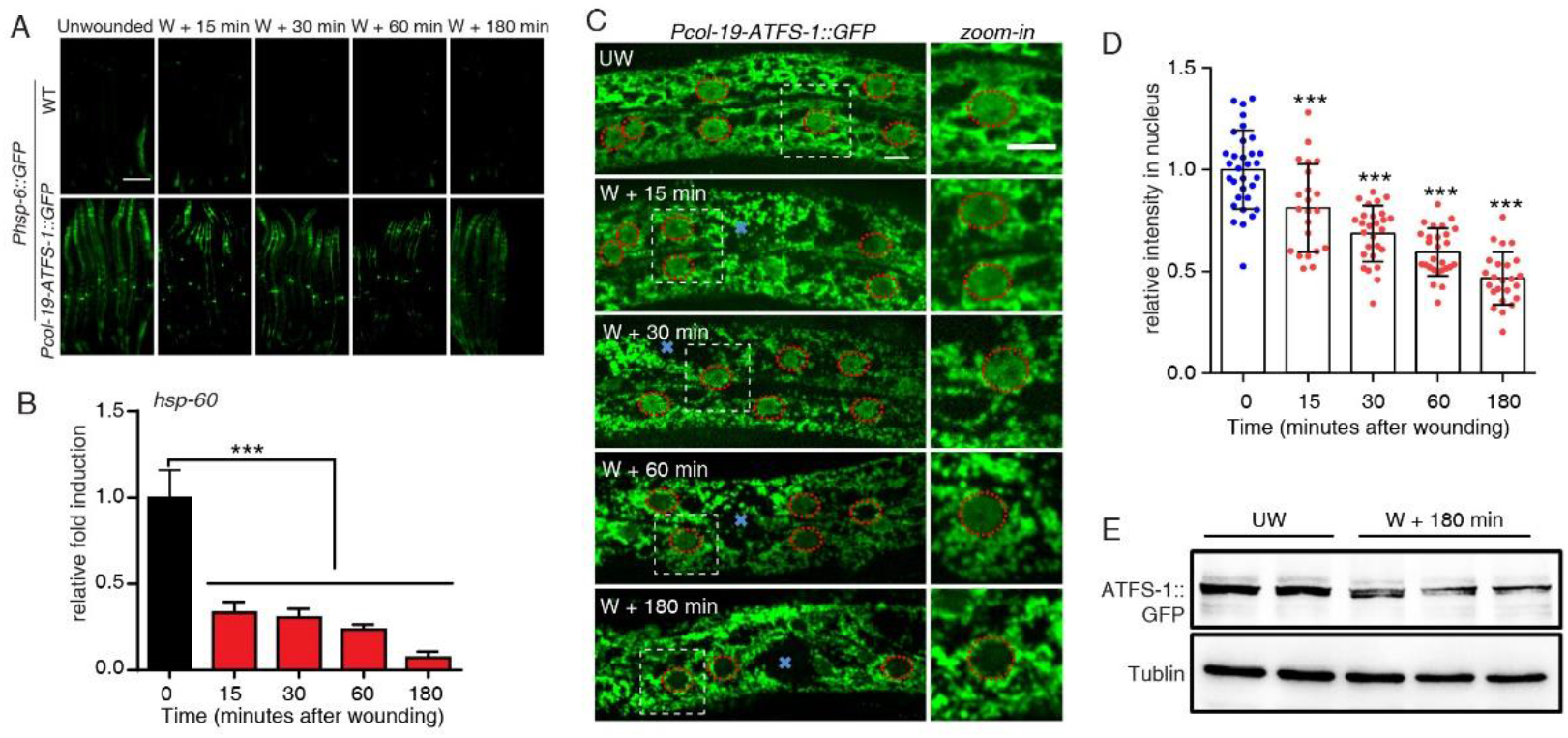
Wounding reduces mitoUPR by inhibition of nuclear localization of ATFS-1. (A)Representative microscopy images of the HSP-6::GFP signal before and at early time points after needle wounding in both WT and *Pcol-19-ATFS-1::GFP(juSi234)* transgenic animals. Scale bars, 200 μm. (B)*hsp-60* expression levels as determined by qPCR in WT animals. 6-8 stabs throughout the whole hypodermis were performed on each animal and 30 D1 animals were collected for each RNA sample. N = 3 replicates. Bars indicate the mean ± SEM. ***, P < 0.001 (Student’s t-test). (C) Representative confocal microscopy images of ATFS-1::GFP before and at different time points after needle wounding. *Pcol-19-ATFS-1::GFP(juSi234)* single-copy insertion transgenic animals were used for wounding and imaging. Animals were wounded with one anterior stab and one posterior stab. Scale bars, 10 μm. (D) Quantification of ATFS-1::GFP fluorescence in the nucleus post wounding. N ≥ 20 animals. Bars indicate the mean ± SEM. ***, P < 0.001, One-way ANOVA and Bartlett’s test. (E) Western blot analysis of ATFS-1::GFP protein levels before and 3 hours after needle wounding. Protein samples were prepared from the total lysate and detected using an anti-GFP antibody.

To further determine how wounding triggers the reduction of mitoUPR, we examined the localization of ATFS-1::GFP before and after wounding. Single-copy insertion of ATFS-1::GFP was localized in both nuclei and mitochondria in the epidermis before wounding (Figure 3C, Figure S4C). Interestingly, the ATFS-1::GFP signal in the nucleus was significantly reduced, starting from 15 minutes after wounding (Figure 3C, D, Figure S4C, D). The ATFS-1::GFP signal almost diminished from the nucleus, but not from mitochondria, 3 hours after wounding (Figure 3C, D, Figure S4C, D). To confirm this observation, we performed western blotting analysis to examine the ATFS-1::GFP protein level and detected a reduction of overall ATFS-1::GFP protein after wounding (Figure 3E). In contrast, we did not observe a significant change of DVE-1::GFP, a transcription factor that acts during stress to induce mitoUPR after wounding (Figure S4E, F). Collectively, these results suggest that epidermal wounding downregulated the nuclear localization of ATFS-1 to reduce the level of mitoUPR.

### mitoUPR plays upstream of p38 MAPK and TGF-β pathways to regulate innate immune response

To determine how mitoUPR regulates wounding-induced innate immune response, we performed a genetic analysis of *atfs-1* with *nsy-1 and sek-1*, which encode key components of the p38 MAP kinase pathway. Loss-of-function mutants *nsy-1(ok593)* and *sek-1(km4)* abolished the p38 MAP kinase cascade (Kim et al., 2002; Pujol et al., 2008), exhibited significant reductions in the extent of *nlp-29* and *nlp-30* induction after wounding (Figure 4A-4D, Figure S5A). Loss-of-function of *atfs-1(tm4525)* enhanced the expression of NLPs in WT but not in *nsy-1(ok593)* or *sek-1(km4)* mutants. In addition, *atfs-1(tm4525)* did not suppress the expression of *tir-1, nsy-1, pmk-1, sek-1* (Figure S5B-S5E). Collectively, these results suggest that mitoUPR signaling plays upstream of the p38 MAP Kinase pathway in regulating NLPs expression.

**Figure 4.**
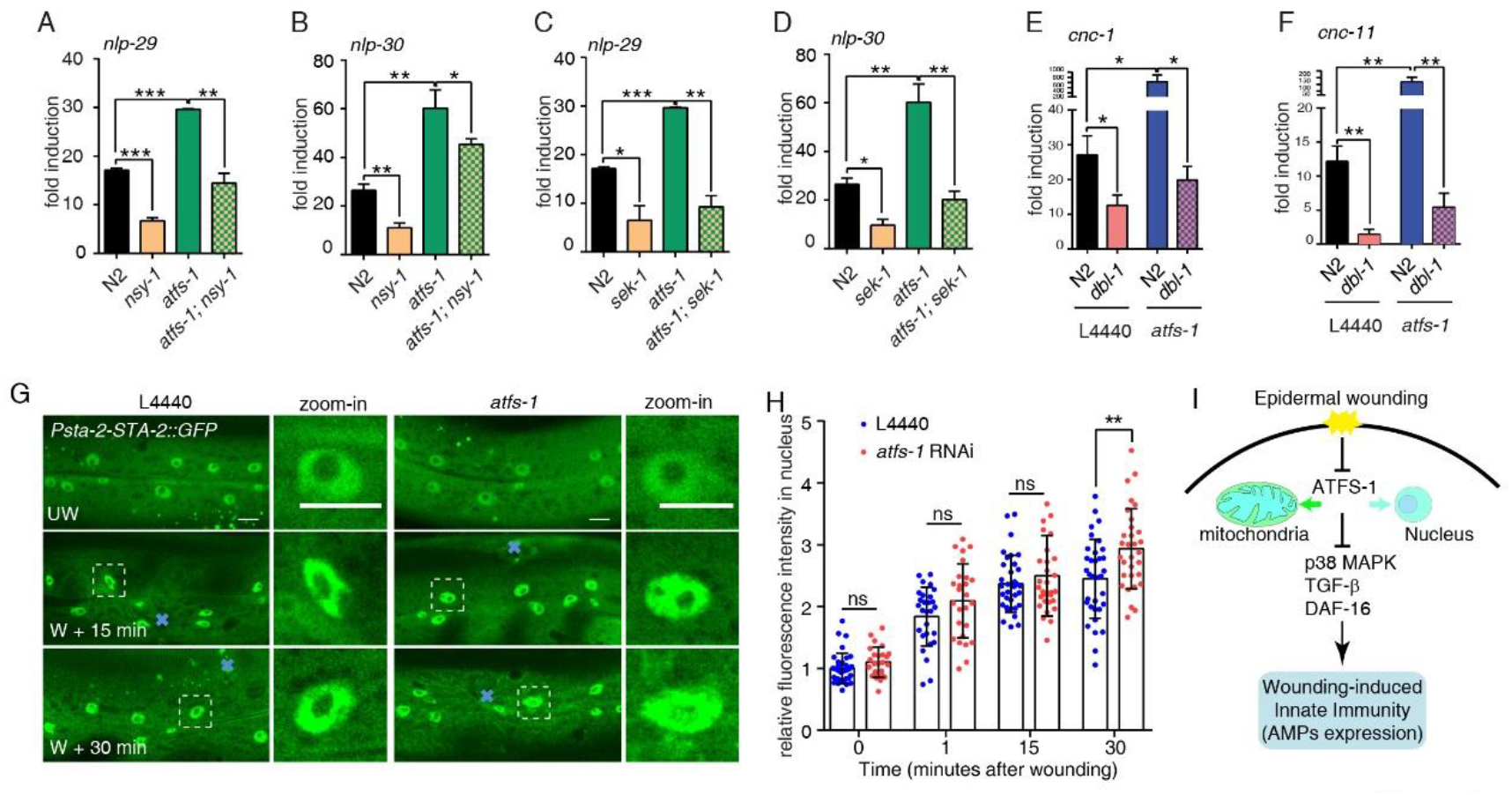
mitoUPR plays upstream of p38MAPK and TGF-β pathways in regulating innate immune response upon wounding. (A-D) *nlp-29* and *nlp-30* expression levels as determined by qPCR in WT, *atfs-1, nsy-1, sek-1*, and double mutants *atfs-1; nsy-1* and *atfs-1; sek-1*. N = 3 replicates. Bars indicate the mean ± SEM. *, P < 0.05, **, P < 0.01, *** P < 0.001 (Student’s t-test). (E-F) *cnc-1* and *cnc-11* expression levels as determined by qPCR in WT or *dbl-1* on control versus *atfs-1*(RNAi). N = 3 replicates. Bars indicate the mean ± SEM. *, P < 0.05, **, P < 0.01 (Student’s t-test). (G) Representative confocal microscopy images of STA-2::GFP fluorescence before and after needle wounding. *Psta-2-STA-2::GFP* transgenic animals were used for wounding and imaging. Scale bars, 10 μm. (H) Quantification of STA-2::GFP fluorescence intensity in the nucleus before and post wounding after feeding with an L4440 empty vector or *atfs-1* dsRNA. N ≥ 20 animals. Bars indicate the mean ± SEM. ns, P > 0.05, **, P < 0.01, One-way ANOVA and Bartlett’s test. (I) A proposed model for mitoUPR senses the epidermal damage and regulates the wounding-induced innate immune response by inhibiting known innate immunue response pathway upon wounding.

Epidermal wounding-induced expression of CNCs is known to be dependent on DBL-1 TGF-β signal pathways (Zugasti and Ewbank, 2009). Similar to the epistatic experiment with the p38 MAP kinase pathway, we observed that *dbl-1(ok3749)* mutation suppressed *cnc-1* and *cnc-11* induction both in WT and *atfs-1(tm4525)* mutant after wounding (Figure 4E-4F, Figure S5A). This result indicates that mitoUPR signaling functions upstream of the TGF-β pathway to regulate the wounding-induced CNCs expression.

Previous work has demonstrated that the p38 MAP kinase cascade activates the STAT-like transcription factor STA-2 to induce an innate immune response (Dierking et al., 2011) and showed that abrogation of STA-2 blocked the defense response to infection and epidermal damage (Dierking et al., 2011; Zhang et al., 2015). To determine how ATFS-1 affects p38 MAP kinase cascade, we examined whether ATFS-1 changes the nuclear localization of the transcription factor STA-2. STA-2::GFP was localized to both cytosolic and nucleus in the unwounded epidermis. However, strikingly, the STA-2::GFP signal intensity was significantly increased in the nucleus as early as 15 minutes after wounding (Figure 4G, H), suggesting that epidermal wounding triggers rapid STA-2 translocation. The loss-of-function of *atfs-1(tm4525)* increased the STA-2::GFP signal in the nucleus within 30 minutes after wounding (Figure 4G, H), indicating that ATFS-1 may inhibit STA-2 translocation in the nucleus and negatively regulates the induction of AMPs after epidermal damage.

## Discussion

In conclusion, our results suggest wounding induces downregulation of mitoUPR to regulate the innate immune response in *C. elegans* epidermis. ATFS-1 and mitoUPR play upstream of well-known p38 MAP kinase and TGF-β pathways to regulate the expression of AMPs (Figure 4I). Mitochondria are protective organelles that have evolved basic physiological processes to maintain homeostasis and defend against environmental insults (Banoth and Cassel, 2018). Our results indicate a previously unknown injury response process in the epidermis that reduces the mitoUPR level, which in turn supports the fine-tuning of the regulation of multiple antimicrobial genes; this process acts upstream of known innate immune pathways upon epidermal wounding.

Stress and infection usually cause increased mitoUPR, a process in which key chaperone proteins clear misfolded and unfolded proteins (Banoth and Cassel, 2018; Lin and Haynes, 2016; Taffoni and Pujol, 2015). One possible explanation for these apparently contradictory conclusions from infection and wounding is that there are no dedicated immune cells in *C. elegans* (Kim and Ewbank, 2018), so specific forms of tissue damage may engage innate immune responses via distinct strategies. Or there could be separate signaling cascades for controlling gene expression after tissue damage vs. pathogen infection by secreting similar antimicrobial peptide genes (Pellegrino et al., 2014; Pujol et al., 2008).

Although the mitoUPR is known to regulate innate immune responses in both the intestine and the epidermis, the signal for induction of mitroUPR might be identified by different receptors and transduced by specific pathways (Nargund et al., 2015; Pellegrino et al., 2014). Furthermore, whether the intestinal endothelial tissue can exhibit a similar immune phenotype as the epidermis upon wounding remains to be determined. Likewise, it is conceivable that the skin has evolved a unique host defense mechanism since it also acts as a first barrier to the surrounding environment in metazoans.

## Materials and Methods

### Strains and transgenic animals

Strains of *C. elegans* were cultured on the nematode growth medium (NGM) plates seeded with *E. coli* OP50 following standard protocols at 20°C(Brenner, 1974). The N2 Bristol strain was used as the wild type (WT) strain. Standard microinjection methods were used to generate transgenic animals carrying extrachromosomal arrays (*zjuEx*). Single-copy insertion of arrays (*zjuSi*) was accomplished by CRISPR-Cas9 mediated gene editing. Strains that were used in this work are summarized in the supplementary Table 1.

### Needle wounding

NGM plates were cooled down on the ice for 20 minutes and transferred 30-40 young adult animals onto each plate. Needle wounding was then performed by microinjection needle on the hyp7 as described previously (Xu and Chisholm, 2011).

### RNAi

The dsRNA bacteria were recovered from -80°C and incubated on the Ampicillin-resistant L.B. plate at 37°C overnight. The next day, the single clone was picked into 1ml Ampicillin-resistant L.B. liquid and incubated for about 6 hours. When the OD reached 0.4-0.6, we sed the bacteria solution on prepared RNAi plates and kept them in the dark. 72 hours later, we prepared a plate of adult animals and collected eggs by bleaching. We then put fresh eggs on the RNAi plate with specific dsRNA and waited for them to grow until L4 at 20°C. We then transferred about 200 synchronized late-L4 animals to a new RNAi plate and collected 30 D1 animals for each RNA sample. The *atfs-1* CDS dsRNA clone was made by PCR from the cDNA.

### RNA isolation and real-time RT–PCR

Worms were plated onto NGM plates and incubated at 20°C. Synchronized late -L4 animals were then transferred to a new plate. Next day, 30 young adult animals for each RNA sample were collected and total RNA was obtained in 500μl Trizol reagent (Invitrogen, Carlsbad, CA, USA). cDNA was then synthesized via the cDNA Synthesis Kit (TOYOBO TRT-101). Using *act-1* as the internal control to normalize expression and determine differences, qRT–PCR was performed according to the SYBR green kit instructions (Vazyme, China). Primer sequences used for qRT–PCR were listed.

### Western blot

For *Pcol-19-ATFS-1::GFP* single-copy insertion transgenic animals, we collected 300 synchronized D1 animals for each protein sample and got the total cell lysate for detection. Before detection, the protein samples were normalized to the same concentration. Tublin was used as the internal control. Western blot was performed according to the Bio-Rad system.

### Imaging analysis

All the images were analyzed using ImageJ software (https://imagej.nih.gov/ij/).

### Statistical analysis

All statistical analyses used GraphPad Prism (La Jolla, CA). Two-way comparisons used Student’s t-test and multiple comparisons used one-way ANOVA Bartlett’s test.

## Acknowledgments

We thank Drs. Xiaochen Wang for reagents. Several *C. elegans* strains were provided by the CGC, which is supported by the NIH Office of Research Infrastructure Programs (P40OD010440). This work is supported by the National Natural Science Foundation of China (31671522, 31972891) to S.X.

## Author contributions

S.X. and J.Z. conceived the study and designed experiments. J.Z. designed and performed the experiments. H.Z. made constructs, performed wounding assay, X.R. performed Western Blot assay. S.X. and J.Z. wrote the manuscript.

## Competing interests

The authors declare no competing financial interests.

## Supplemental Information

Including 5 Supplementary Figures and Figure Legends

**Figure 1.**
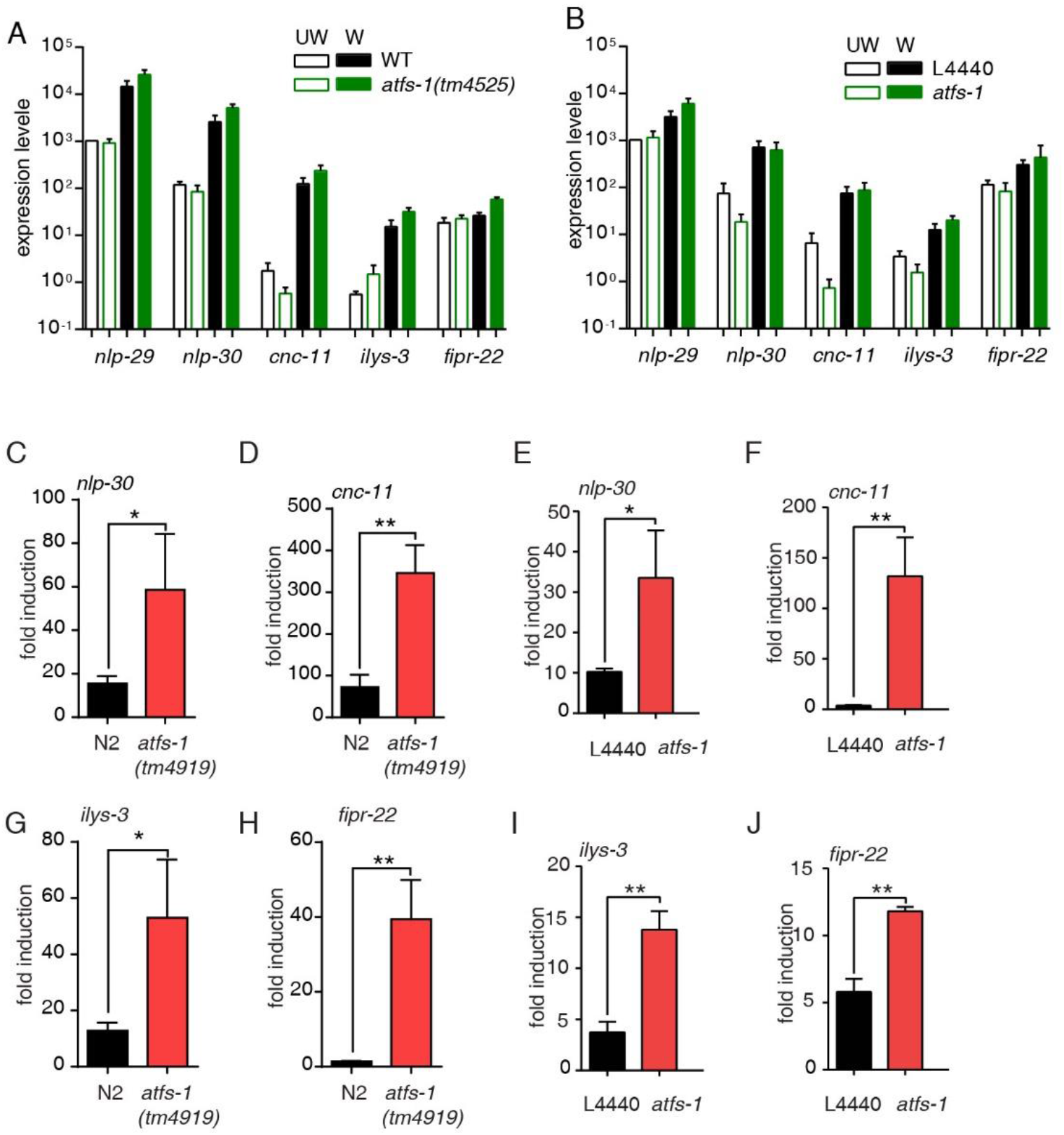
Loss-of-function of ATFS-1 enhances the epidermal wounding-induced innate immune response in *C. elegans* epidermis. (A) Relative expression of *nlp-29, nlp-30, cnc-1, cnc-4, ilys-3* and *fipr-22* as determined by qRT–PCR in the wounded animals in both wild type (WT) and atfs- 1(tm4525) mutant. N = 3. Bars indicate mean ± SEM. (B) Relative expression of *nlp-29, nlp-30, cnc-1, cnc-4, ilys-3* and *fipr-22* as determined by qRT–PCR in the wounded animals in both atfs-1 RNAi and control L4440 (empty vector) RNAi animals. N = 3. Bars indicate mean ± SEM. (C-J) Fold ineduction analysis of *nlp-30, cnc-11, ilys-3* and *fipr-22* transcripts as determined by qRT–PCR in WT and *atfs-1(4919)* or *atfs-1* RNAi animals. N = 3. Bars indicate mean ± SEM. *, P < 0.05, **, P < 0.01, ***, P < 0.001 (student’s t-test).

**Figure 2.**
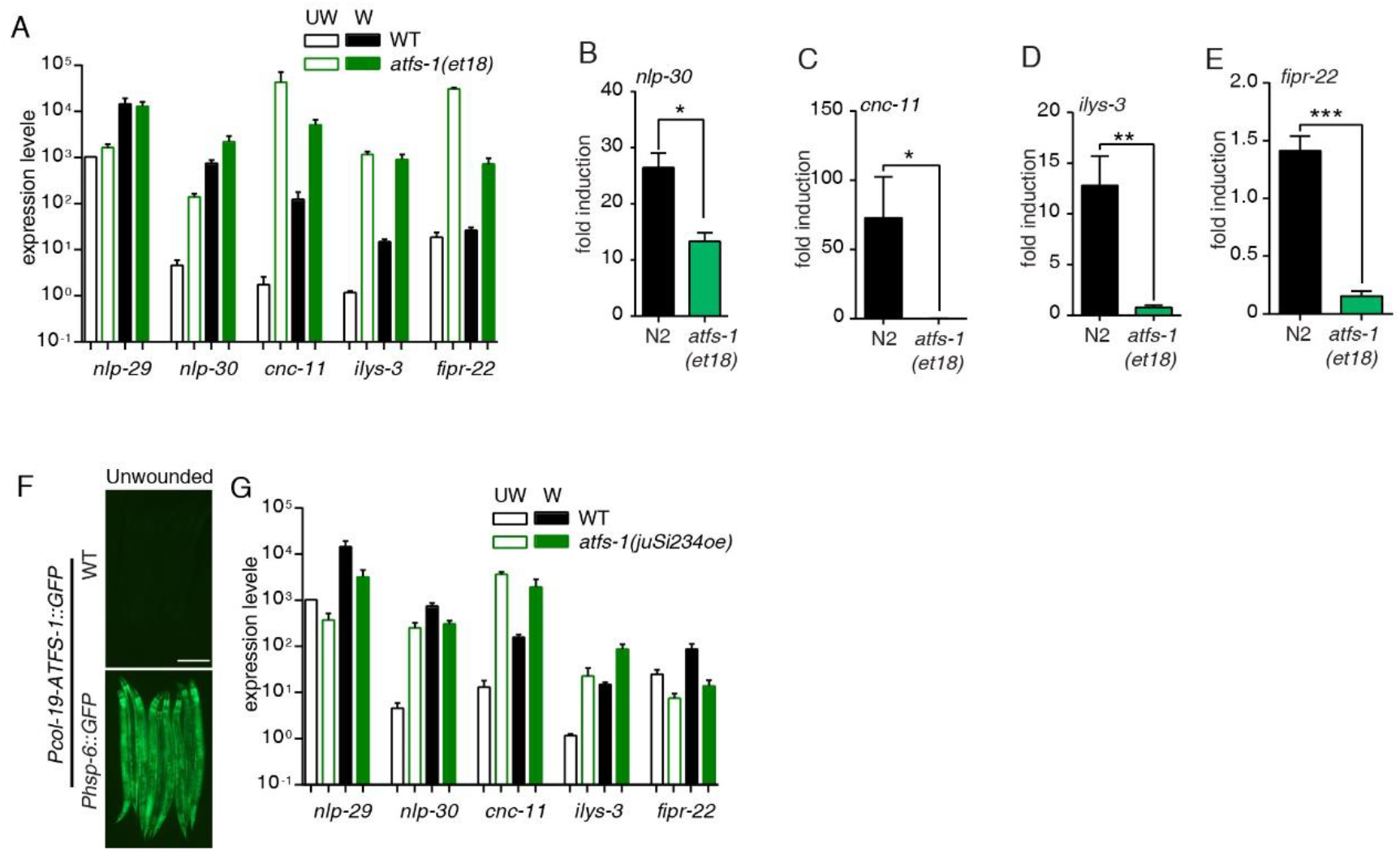
Elevated mitoUPR inhibits the induction of wounding-induced innate immune response genes. (A) Relative expression of *nlp-29, nlp-30, cnc-1, cnc-4, ilys-3* and *fipr-22* as determined by qRT–PCR in the wounded animals in both WT and *atfs-1(et18gf)* mutant. N = 3. Bars indicate mean ± SEM. (B-E) *nlp-30, cnc-11, ilys-3,and fipr-22* transcripts as determined by qRT–PCR in both WT and *atfs-1(et18gf)* mutants. N = 3. Bars indicate mean ± SEM. *, P < 0.05, **, P < 0.01 (student’s t-test). (F) The fluorescence signal of *Phsp-6::GFP* in both WT and *Pcol-19-ATFS- 1::GFP(juSi234)* overexpressed animals. Scale bars, 200 μm. (G) Relative expression of *nlp-29, nlp-30, cnc-1, cnc-4, ilys-3* and *fipr-22* as determined by qRT–PCR in the wounded animals in both WT and *atfs-1(juSi234)* overexpressed animals. N = 3. Bars indicate mean ± SEM.

**Figure 3.**
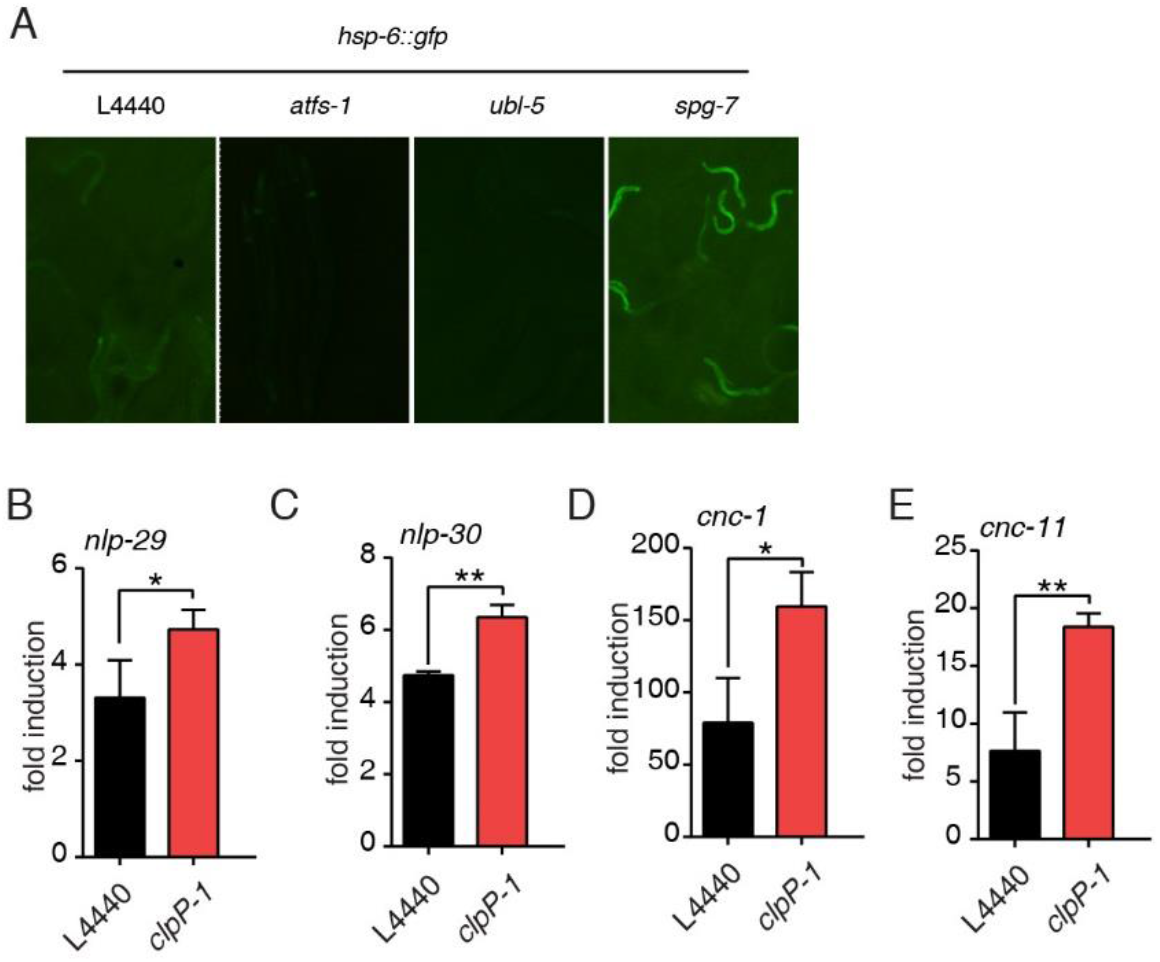
Low level of mitoUPR enhance the induction of AMPs after wounding. (A) Representative microscopy images of the HSP-6::GFP signal in the unwounded animals fed with an L4440 empty vector, *atfs-1, spg-7*, or *ubl-5* dsRNA. Scale bars, 500 μm. *(B) nlp-29, nlp-30, cnc-1* and *cnc-11* transcripts as determined by qRT–PCR in the wounded animals fed by L4440 empty vector or *clpP-1* dsRNA. N = 3. Bars indicate mean ± SEM. *, P < 0.05, **, P < 0.01 (student’s t-test).

**Figure 2.**
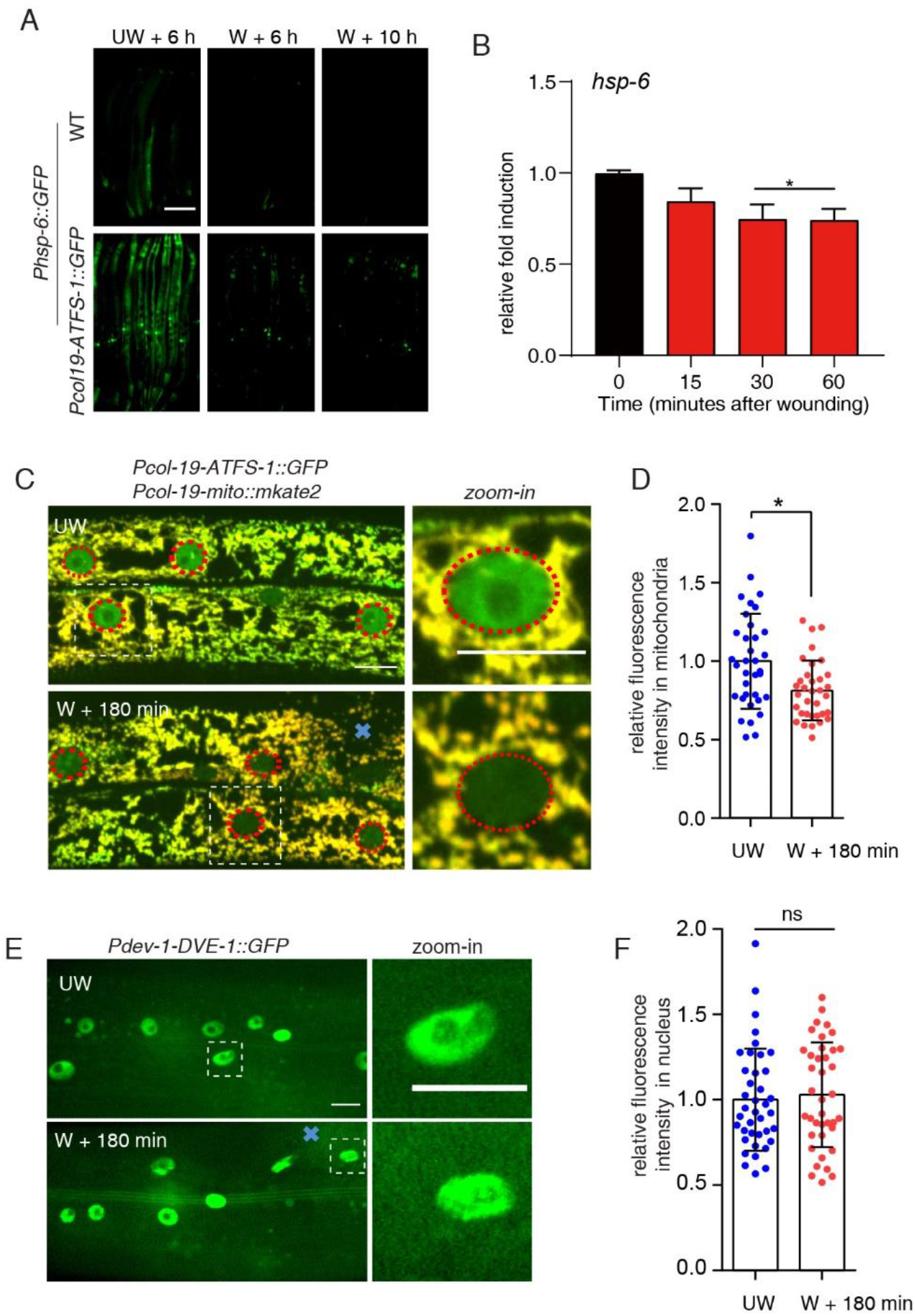
Epidermal wounding inhibits ATFS-1-dependent UPR^mt^. (A) Representative microscopy images of HSP-6::GFP signal before and at later times after needle wounding in both wild-type and *Pcol-19-ATFS-1::GFP(juSi234)* transgenic animals. HSP-6::GFP signal became weaker following wounding. Scale bars, 200 μm. *(B) hsp-6* expression levels as determined by qPCR in WT animals. 6-8 stabs throughout the whole hypodermis were performed on each animal and 30 young adult animals were collected for each RNA sample. N = 3 replicates. Bars indicate the mean ± SEM. ***, P < 0.001 (Student’s t-test). (C) Representative confocal microscopy images of ATFS-1::GFP and mito::mKate2 before and 3 hours post wounding. *Pcol-19-ATFS-1::GFP(juSi234); Pcol-19- mito::mKate2(zjuSi47)* transgenic animals were used for wounding and imaging. Scale bars, 10μm. (D) Statistic quantification of ATFS-1::GFP fluorescence intensity in mitochondria 3 hours post wounding. *Pcol-19-ATFS-1::GFP(juSi234);Pcol-19- mito::mKate2(zjuSi47)* transgenic animals were used for wounding and statistics. The result was the ratio of green fluorescence to red fluorescence. N ≥ 20. Bars indicate mean ± SEM. *, P<0.05 (Student’s t-test). (E) Representative confocal microscopy images of DVE-1::GFP in nucleus before and 3 hours post wounding. *Pdve-1-DVE-1::GFP(zcIs39)* transgenic animals were used for wounding and imaging. Scale bars, 10μm. (F) Statistic quantification of DVE-1::GFP fluorescence intensity in nucleus. N ≥ 20. Bars indicate mean ± SEM. ns, P > 0.05 (student’s t-test).

**Figure 5.**
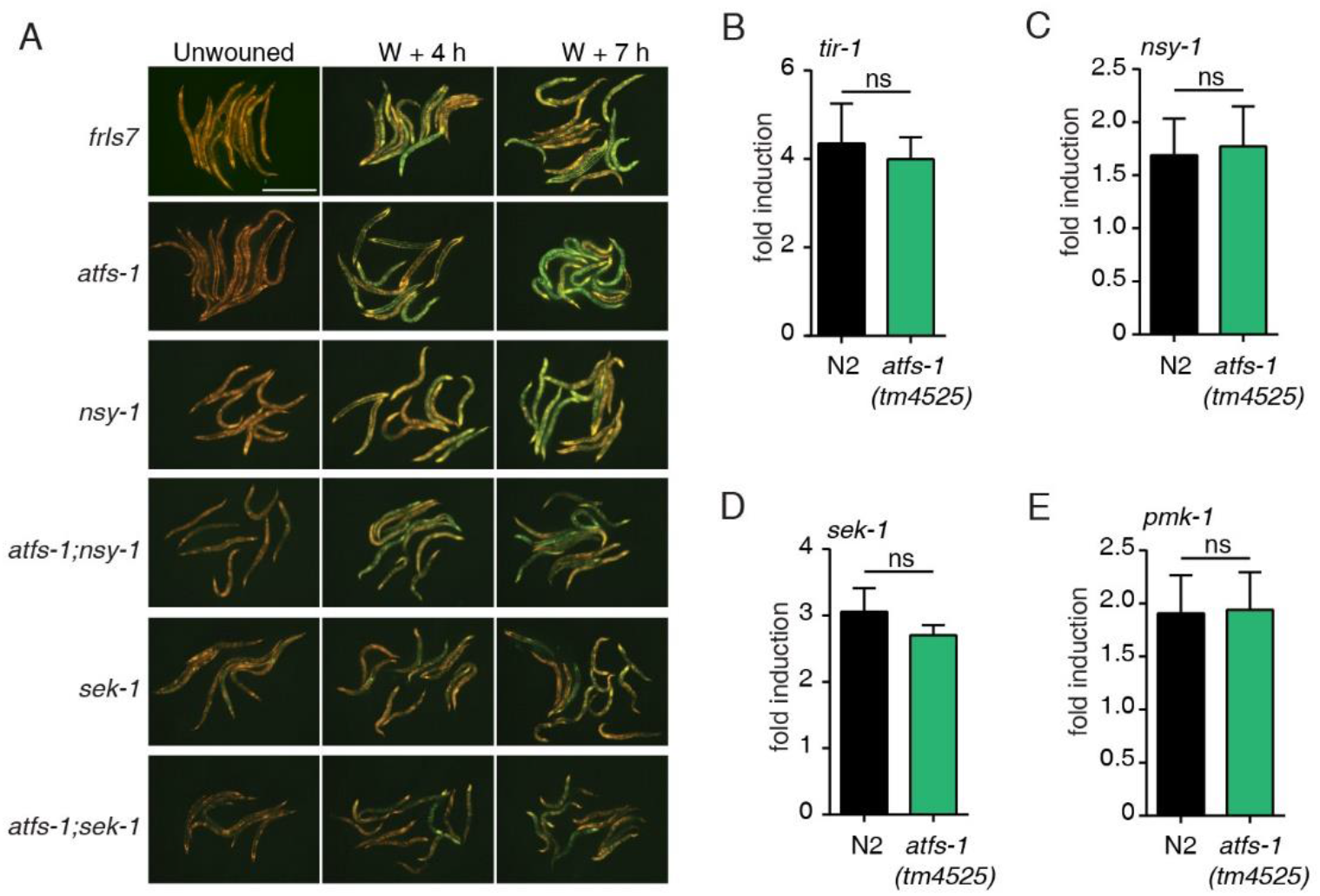
ATFS-1 acts upstream of p38MAP kinase and TGF-b pathway. (A)The green fluorescence in transgenic worms carrying a *Pnlp-29::GFP* reporter was observed post wounding in WT, *atfs-1, nsy-1, sek-1* and double mutants *atfs-1; nsy-1* and *atfs-1; sek-1*. Scale bars, 500mm. (B-E) *tir-1, nsy-1, sek-1* and *pmk-1* transcripts as determined by qRT–PCR in WT and *atfs-1(tm4525)*. N = 3. Bars indicate mean ± SEM. ns, P > 0.05 (student’s t-test).

## Notes

### Competing Interest Statement

The authors have declared no competing interest.

## References

Banoth, B., and Cassel, S.L. (2018). Mitochondria in innate immune signaling\. Transl Res 202, 52–68.

Benedetti, C., Haynes, C.M., Yang, Y., Harding, H.P., and Ron, D. (2006). Ubiquitin-like protein 5 positively regulates chaperone gene expression in the mitochondrial unfolded protein response. Genetics 174, 229–239.

Brenner, S. (1974). The genetics of Caenorhabditis elegans. Genetics 77, 71–94.

Dierking, K., Polanowska, J., Omi, S., Engelmann, I., Gut, M., Lembo, F., Ewbank, J.J., and Pujol, N. (2011). Unusual regulation of a STAT protein by an SLC6 family transporter in C. elegans epidermal innate immunity. Cell Host Microbe 9, 425–435.

Dunbar, T.L., Yan, Z., Balla, K.M., Smelkinson, M.G., and Troemel, E.R. (2012). C. elegans detects pathogen-induced translational inhibition to activate immune signaling. Cell Host Microbe 11, 375–386.

Gravato-Nobre, M.J., Vaz, F., Filipe, S., Chalmers, R., and Hodgkin, J. (2016). The Invertebrate Lysozyme Effector ILYS-3 Is Systemically Activated in Response to Danger Signals and Confers Antimicrobial Protection in C. elegans. PLoS Pathog 12, e1005826.

Haynes, C.M., Petrova, K., Benedetti, C., Yang, Y., and Ron, D. (2007a). ClpP mediates activation of a mitochondrial unfolded protein response in C. elegans. Dev Cell 13, 467–480.

Haynes, C.M., Petrova, K., Benedetti, C., Yang, Y., and Ron, D. (2007b). ClpP mediates activation of a mitochondrial unfolded protein response in C. elegans. Dev Cell 13, 467–480.

Haynes, C.M., and Ron, D. (2010). The mitochondrial UPR - protecting organelle protein homeostasis. J Cell Sci 123, 3849–3855.

Kim, D.H., and Ewbank, J.J. (2018). Signaling in the innate immune response. WormBook 2018, 1–35.

Kim, D.H., Feinbaum, R., Alloing, G., Emerson, F.E., Garsin, D.A., Inoue, H., Tanaka-Hino, M., Hisamoto, N., Matsumoto, K., Tan, M.W., et al. s(2002). A conserved p38 MAP kinase pathway in Caenorhabditis elegans innate immunity. Science 297, 623–626.

Lin, Y.F., and Haynes, C.M. (2016). Metabolism and the UPR(mt). Mol Cell 61, 677–682.

Liu, Y., Samuel, B.S., Breen, P.C., and Ruvkun, G. (2014). Caenorhabditis elegans pathways that surveil and defend mitochondria. Nature 508, 406–410.

Melo, J.A., and Ruvkun, G. (2012). Inactivation of conserved C. elegans genes engages pathogen-and xenobiotic-associated defenses. Cell 149, 452–466.

Nargund, A.M., Fiorese, C.J., Pellegrino, M.W., Deng, P., and Haynes, C.M. (2015). Mitochondrial and nuclear accumulation of the transcription factor ATFS-1 promotes OXPHOS recovery during the UPR(mt). Mol Cell 58, 123–133.

Nargund, A.M., Pellegrino, M.W., Fiorese, C.J., Baker, B.M., and Haynes, C.M. (2012). Mitochondrial import efficiency of ATFS-1 regulates mitochondrial UPR activation. Science 337, 587–590.

Pellegrino, M.W., Nargund, A.M., Kirienko, N.V., Gillis, R., Fiorese, C.J., and Haynes, C.M. (2014). Mitochondrial UPR-regulated innate immunity provides resistance to pathogen infection. Nature 516, 414–417.

Pujol, N., Cypowyj, S., Ziegler, K., Millet, A., Astrain, A., Goncharov, A., Jin, Y., Chisholm, A.D., and Ewbank, J.J. (2008). Distinct innate immune responses to infection and wounding in the C. elegans epidermis. Curr Biol 18, 481–489.

Rauthan, M., Ranji, P., Aguilera Pradenas, N., Pitot, C., and Pilon, M. (2013). The mitochondrial unfolded protein response activator ATFS-1 protects cells from inhibition of the mevalonate pathway. Proc Natl Acad Sci U S A 110, 5981–5986.

Richardson, C.E., Kooistra, T., and Kim, D.H. (2010). An essential role for XBP-1 in host protection against immune activation in C. elegans. Nature 463, 1092–1095.

Taffoni, C., and Pujol, N. (2015). Mechanisms of innate immunity in C. elegans epidermis. Tissue Barriers 3, e1078432.

Xu, S., and Chisholm, A.D. (2011). A Galphaq-Ca(2)(+) signaling pathway promotes actin-mediated epidermal wound closure in C. elegans. Curr Biol 21, 1960–1967.

Yoneda, T., Benedetti, C., Urano, F., Clark, S.G., Harding, H.P., and Ron, D. (2004). Compartment-specific perturbation of protein handling activates genes encoding mitochondrial chaperones. J Cell Sci 117, 4055–4066.

Zhang, Y., Li, W., Li, L., Li, Y., Fu, R., Zhu, Y., Li, J., Zhou, Y., Xiong, S., and Zhang, H. (2015). Structural damage in the C. elegans epidermis causes release of STA-2 and induction of an innate immune response. Immunity 42, 309–320.

Zugasti, O., and Ewbank, J.J. (2009). Neuroimmune regulation of antimicrobial peptide expression by a noncanonical TGF-beta signaling pathway in Caenorhabditis elegans epidermis. Nat Immunol 10, 249–256.

